# WntA expression and wing transcriptomics illuminate the evolution of stripe patterns in skipper butterflies

**DOI:** 10.1101/2025.09.28.679064

**Authors:** Jasmine D. Alqassar, Teomie S. Rivera-Miranda, Joseph J. Hanly, Christopher R. Day, Silvia M. Planas Soto-Navarro, Paul B. Frandsen, Riccardo Papa, Arnaud Martin

**Affiliations:** Department of Biological Sciences, The George Washington University, Washington, DC, USA; Department of Biology, University of Puerto Rico-Río Piedras, San Juan, Puerto Rico; Smithsonian Tropical Research Institute, Gamboa, Panama; Molecular and Cellular Biology Laboratory, National Institute of Environmental Health Sciences, Durham, USA; Molecular Sciences and Research Center, University of Puerto Rico-Río Piedras, San Juan, Puerto Rico; Department of Plant and Wildlife Sciences, Brigham Young University, Provo, UT, USA; Dipartimento di Scienze Chimiche della Vita e della Sostenibilità Ambientale, Università di Parma, Italy

**Keywords:** evo-devo, pattern formation, wing epithelium, WntA signaling, Hesperiidae, Nymphalid Ground Plan, developmental homology, proximo-distal patterning

## Abstract

Skippers (Hesperiidae) form a distinct lineage of butterflies where the developmental mechanisms of color patterning have seldom been studied. The wing patterns of skippers often consist of median stripes, and classic comparative morphology studies suggest these elements are homologous to the Central Symmetry System (CSS) found in nymphalid butterflies. Consistent with this hypothesis, we show here that expression of the signaling ligand gene *WntA*, a marker of the CSS in nymphalids, prefigures the position of the main wing patterns of the silver-spotted skipper *Epargyreus clarus*. Notably, *WntA* expression is associated with different color outputs in the forewing vs. hindwing, validating a theory of pattern homology from the mid-twentieth century. To further gain insights into the genes associated with this patterning process, we generated an annotated genome for *E. clarus* and performed an RNAseq study profiling gene expression along the proximo-distal (P-D) axis of early pupal wings. These data suggest that the transcription factor genes *lobe, u-shaped*, and *odd-paired* are expressed in restricted P-D sections of the wing similarly to *WntA*, indicating they may participate in the patterning of the CSS. In addition, inverted expression patterns of *dachsous* and *four-jointed*, as well as the expression of transcription factors with roles in specifying proximal (*homothorax, tiptop*/*teashirt*) and distal (*vestigial, scalloped*) section of ths *Drosophila* wing disk, suggest a deep conservation of wing P-D patterning process between Diptera and Lepidoptera. This work highlights the developmental homology between the CSS of Hesperiidae and Nymphalidae, and enriches our knowledge of patterning mechanisms in lepidopteran wings.

**Summary Statement:** This study reveals how skipper butterfly wing stripes share deep developmental origins with other butterflies, offering fresh insight into wing pattern evolution.

## Introduction

How do butterflies make color patterns on their wings, and to what extent can these various stripes and spots be considered homologous across large phylogenetic distances? The Hesperiidae family includes at least 3,800 species of skipper butterflies (Bridges, 1994; Van Nieukerken et al., 2011) and is uniquely situated to help answer this question. Hesperiidae diverged about 95 MYA from the core lineage that encompasses the main butterfly families (Kawahara et al., 2023), including Nymphalidae which represents the main model systems for studying developmental evolution. Skippers show dazzling variation in their wing patterns, and while the interest in their diversification was recently reignited by phylogenetic and population genomics studies (Li et al., 2019; Toussaint et al., 2018; Toussaint et al., 2025), few studies have investigated its developmental basis.

Deciphering the degrees of homology between butterfly wing patterns is crucial to understanding whether conserved processes and mechanisms potentiate their diversification. In the 1920s, comparative morphologists Süffert and Schwanwitsch described the wings of Nymphalidae as deriving from pattern elements known as symmetry systems that together form a homology system known as the nymphalid ground plan (Nijhout, 1978; Nijhout, 1991; Schwanwitsch, 1924; Süffert, 1927). Following the terminology of Schwanwitsch (**Fig. 1A-B**), the Central Symmetry System (CSS) is a prominent feature of this ground plan, taking the form of a color field in the median region of the wing, and delimited on each side by proximal and distal boundaries (called **M_1_** and **M_2_**). Schwanwitsch spent most of his career building a theory of pattern homology across Lepidoptera that extends the existence of putative CSS homologues throughout the entire order (Schwanwitsch, 1956). For instance, this model proposes that Hesperiidae show a CSS element similar to the one found in Nymphalidae, as illustrated by Schwanwitsch in the skipper *Pyrgus sidae* (**Fig. 1C**). Interestingly, if correct, this prediction implies that identifying the hesperiid CSS: *(a)* forewings show a dislocation of the CSS, which is more distal in its anterior half; *(b)* in hindwings, the CSS is often white or reflective, unlike in the forewings. This model makes homology calls that are counterintuitive, because this results in CSS elements that can shift in color within the same individual: in *P. sidae*, the CSS is black and gray in the forewing, and white in the hindwing.

**Figure 1.**
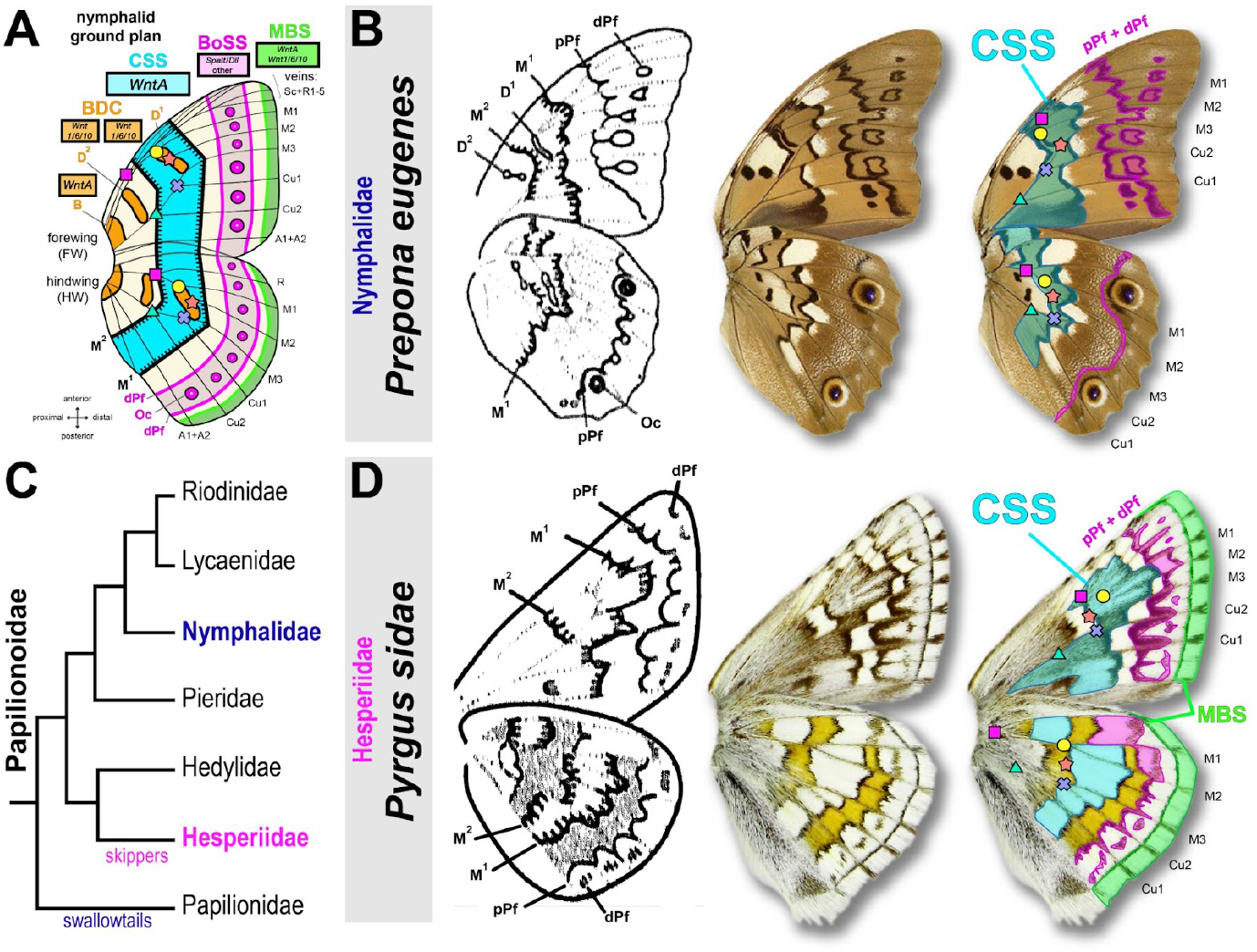
Hypotheses of stripe pattern homology between nymphalids and hesperiids. **A**. Current summary of the nymphalid ground plan, mainly based on the terminology of Schwanwitsch (Nijhout, 1991; Schwanwitsch, 1924) and including recent updates by Otaki and Mazo-Vargas *et al*. (Mazo-Vargas et al., 2017a; Otaki, 2012; Otaki, 2021; Schwanwitsch, 1956). Discalis elements (D_1_ and D_2_); CSS: Central Symmetry System (cyan); BoSS: Border Ocelli Symmetry System (magenta); Oc: forewing Border Ocelli; pPf and dPf: proximal and distal Parafocal elements; MBS: Marginal Band System (green). Colored vignettes denote vein intersection landmarks. Magenta square: junction between R and M vein trunks; yellow dot: M_1_-M_2_ junction; red star: junction between discal crossvein and M_3_; blue cross: M_3_-Cu_2_ junction; green triangle: Cu_1_-Cu_2_ junction. Rectangles feature the name of marker genes. **B**. Ventral wing patterns of the nymphalid *Prepona eugenes* with ground plan annotations proposed by Schwanwitsch, 1956 (left: reproduction of published drawings; right: equivalent annotations as color overlays). **C**. Phylogenetic relationship between Papilionoidae families. **D**. Ventral wing patterns of the hesperiid *Pyrgus sidae* annotated as in panel A, and highlighting the inferred CSS predicted by Schwanwitsch, 1956 (left panel). According to this author, the CSS marks a grey pattern in forewings, and a dislocated white stripe pattern in hindwings, suggesting uncoupling of pattern and color state in fore/hindwings in skippers.

Gene expression provides important clues for probing homology relationships in developmental processes (DiFrisco et al., 2020; Pantalacci and Sémon, 2015). The gene *WntA* has emerged as a reliable spatial marker of the CSS and other patterns in nymphalid developing wing tissues (Concha et al., 2019; Gallant et al., 2014; Martin and Courtier-Orgogozo, 2017; Martin and Reed, 2014a). *WntA* encodes a secreted ligand of the Wnt family that requires the Frizzled2 receptor to induce patterns of the basal, central, and marginal patterning systems (Banerjee et al., 2023; Hanly et al., 2021; Hanly et al., 2023; Mazo-Vargas et al., 2017a; Mazo-Vargas et al., 2022a). WntA signaling has been understudied because this gene was lost both in *Drosophila* and in the vertebrate lineage, and little is known about its modes of signal transduction and transcriptional regulation.

Here we selected the silver-spotted skipper *Epargyreus clarus* as a model system for the study of hesperiid wing pattern, as this North American skipper is easily collected on invasive kudzu patches, where larvae inhabit characteristic leaf shelters (Lind et al., 2001). We generated a genome assembly for this species and profiled the spatial expression of *WntA* to test for the homologies of the hesperiid and nymphalid CSS patterns. Then, we used RNAseq of pupal wing sections to conduct Differential Gene Expression analyses comparing the transcriptomes of proximal, medial, and distal regions of pupal wings. We discuss how *WntA* expression supports the long-standing pattern homology system proposed by Schwanwitsch, and detect a few candidate transcription factors that may be involved in the proximo-distal patterning of these systems in skippers and beyond.

## Results

### A reference genome for Hesperiidae

We generated a reference genome (**Fig. 2A-B**) and transcriptomes for *E. clarus* to enable future studies of Hesperiidae. High-molecular weight DNA from a female pupa was sequenced with PacBio HiFi reads, and assembled into a haplotype of 40 contigs and a total genome size of 452 Mb (N50 = 15.8 Mb; BUSCO analysis scores: Complete Single-copy: 99.4%, Complete Duplicated: 0.4%, Fragmented: 0.00%, Missing: 0.2%), including a complete mitogenome of 30.6 kb (NW_027332243.1). Assignments of the 32 main nuclear contigs to Merian elements (**Fig. 2C-D**), a system of homologous linkage groups conserved across Lepidoptera, recovered the set of 31 chromosomes ancestral to the Ditrysia lineage (Wright et al., 2024), implying the absence of any major chromosomal rearrangement in this species for about 150MY (Kawahara et al., 2019). Two contigs are splitting the M30 element, but are unlikely to represent a real fission event, as they likely account for the discrepancy between our contig count (n = 32) and the expected count of n = 31 chromosomes established from the *E. clarus* karyotype (Maeki, 1961).

**Figure 2.**
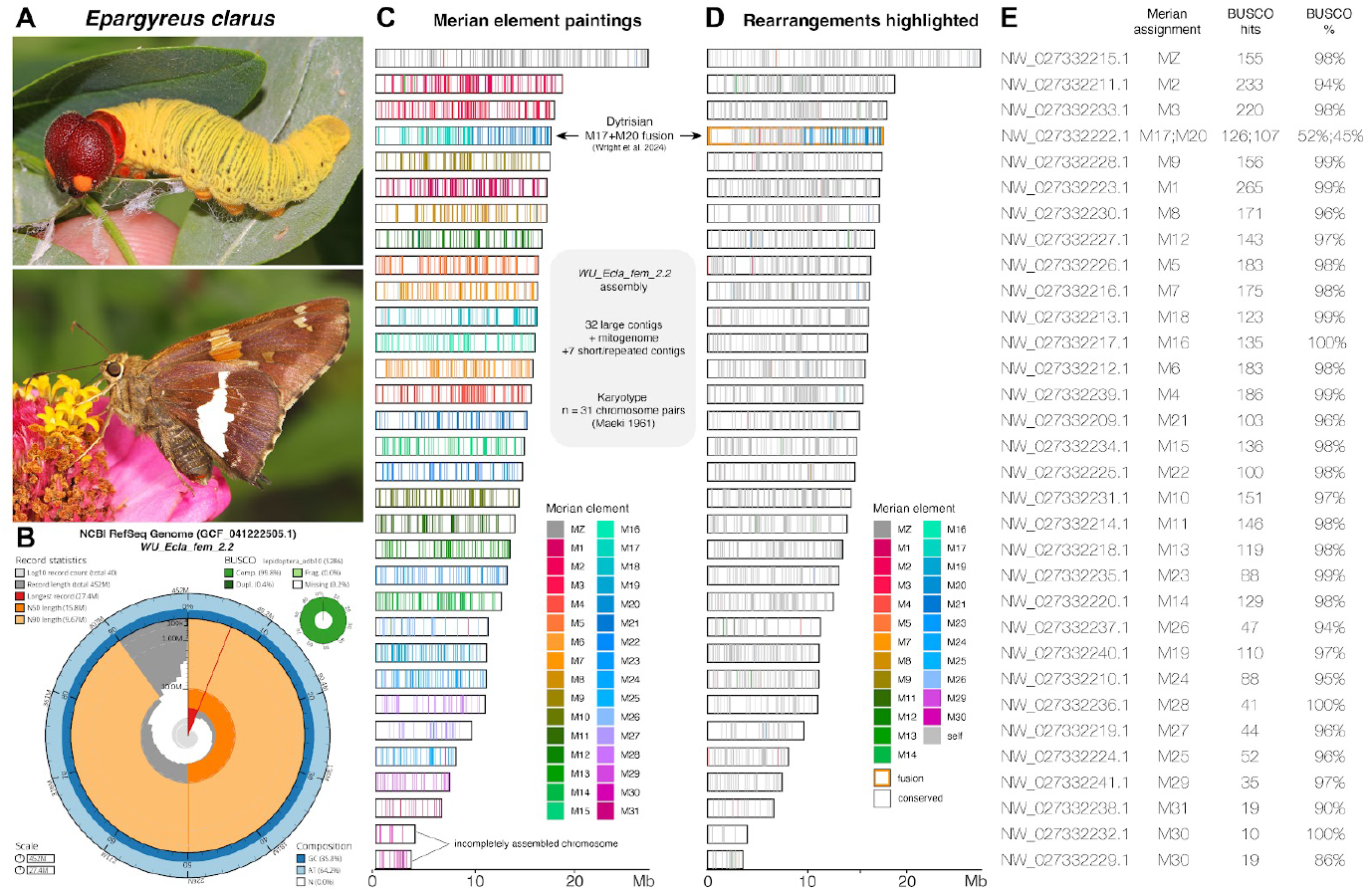
Assignment of the *E. clarus* genome to ancestral Merian elements. **A**. Fifth instar larva and adult form of the Silver-Spotted Skipper *E. clarus* (Photo credits, Judy Gallagher, CC-BY-2.0 license). **B**. Summary SnailPlot assembly and completeness metrics for the *WU_Ecla_fem_2.2* RefSeq genome assembly of *E. clarus* (Challis et al., 2020). **C-D**. Merian elements mapped across chromosomes in the *WU_Ecla_fem_2.2* haploid assembly of *E. clarus*. N = 32 contigs are drawn to scale. BUSCO paintings with orthologue positions are shown as coloured bars, each shaded according to its corresponding Merian element (**C**), or with discrepancies with the ancestral lepidopteran linkage groups highlighted (**D**). The M17-M20 fusion is ancestral to Dytrisia, a lineage that encompasses most of lepidopteran diversity (Kawahara et al., 2019; Wright et al., 2024). **E**. Merian assignments for each *E. clarus* contigs.

Transcriptomes include larval and pupal wings, whole heads from immatures and adults, ovaries, testes, and silk glands, and were generated using poly-dT cDNA synthesis (mRNA sequencing). Two samples were also prepared for total RNA sequencing using a ribosome depletion kit (QIAseq FastSelect –rRNA Fly Kit). This depletion method failed, resulting in over-representation of rRNA, but nonetheless includes non-coding transcripts that may increase transcript diversity.

A genome annotation was generated by the NCBI Eukaryotic Genome Annotation Pipeline team, and encompasses 15,809 predicted gene models, including 13,193 protein-coding genes. This high quality genome assembly and annotation for *E. clarus* are available online at the NCBI Datasets repository (*WU_Ecla_fem_2.2*, GCF_041222505.1) and represent the first NCBI RefSeq genome for Hesperiidae, a diverse family that includes more an estimated 3,800-4,100 skipper species (Bridges, 1994; Van Nieukerken et al., 2011). The *E. clarus* genome encodes the 8 Wnt ligand genes previously found in other lepidopteran genomes (Ding et al., 2019; Fenner et al., 2020; Hanly et al., 2023; Holzem et al., 2019), including *WntA* and the conserved *Wnt1-Wnt9-Wnt6-Wnt10* cluster (**Fig. S1**). Because skippers occupy a key phylogenetic position (**Fig. 1C**) between the core group of butterflies and the early-diverging swallowtail lineage (Espeland et al., 2018; Kawahara et al., 2023), this reference genome fills a gap for comparative genomics in butterflies.

### *WntA* expression and heparin injections unveil the hesperiid CSS

We profiled the spatial expression of *WntA* mRNA using colorimetric *in situ* hybridizations in the fifth instar larval wing disks and 17% pupal wings, two stages where *WntA* marks presumptive color patterns in nymphalids (Banerjee et al., 2023; Concha et al., 2019; Gallant et al., 2014; Hanly et al., 2023; Huber et al., 2015; Martin and Reed, 2014b; Martin et al., 2012; Mazo-Vargas et al., 2017b). Similarly to nymphalids, *E. clarus WntA* is expressed in the medial region of the wing and prefigures the position of adult patterns (**Fig. 3A-B**). In ventral hindwings, *WntA* clearly prefigures the silver bands. In forewings, *WntA* flanks the orange bands and spots and is instead expressed in brown-melanic regions. For example *WntA* expression forms a triangle at the base of the M3-Cu1 vein intersection (blue cross in **Fig. 3**) that corresponds to a melanic region in adults, and that is immediately posterior to a small orange dot. This stereotypical expression of *WntA* at the base of the M1-Cu2 wing compartment resembles the expression of WntA in *Heliconius* and *Limenitis* species: in these nymphalids WntA determines the shape of light-colored medial patterns by inducing melanic patterns that have been called “shutters”, by analogy with shutters than slide onto a window and restrict light (Gallant et al., 2014; Gilbert, 2003; Martin et al., 2012; Nijhout et al., 1990). To test if WntA functionally induces distinct patterns in forewings and hindwings, we initially attempted CRISPR knock-outs but failed to microinject *E. clarus* eggs, due to the unusual toughness of their chorion. Instead, we used heparin injections as previously used in pre-CRISPR studies (Martin and Reed, 2014b; Serfas and Carroll, 2005). Doses of 10-30 μg of heparin, administered within 16 h after pupation, were shown to consistently induce WntA gain-of-function effects across nymphalids, likely because heparin mimics the heparan sulfate proteoglycans of the extracellular matrix that facilitate Wnt transport and uptake (Binari et al., 1997; Gallant et al., 2014; Greco et al., 2001; Mazo-Vargas et al., 2017b;Mazo-Vargas et al., 2022a; Sourakov, 2018; Sourakov and Shirai, 2020). Following injection in the abdomen of early pupae, the forewing orange bands were either reduced at the lower dose (12.5 μg) or completely absent at the higher doses (20-25 μg), an effect that emulates heparin effects in *Heliconius* and *Limenitis* and indicates an expansion of melanic shutter patterns (**Fig. 3C-D**). Conversely, while the hindwing silver band showed a small expansion of its anterior section under lower doses, higher doses resulted in drastic expansion and fuzziness of this pattern (**Fig. 3C-D**). Despite the caveat that heparin injections may affect other signaling ligands, the combination of *WntA* expression assays and pharmacological perturbation strongly indicate that WntA functions as a melanic shutter in forewings and as a silver band activator in hindwings (**Fig. 3E**). These distinct effects reveal the dual nature of the CSS homolog in the hesperiid *E. clarus*.

**Figure 3.**
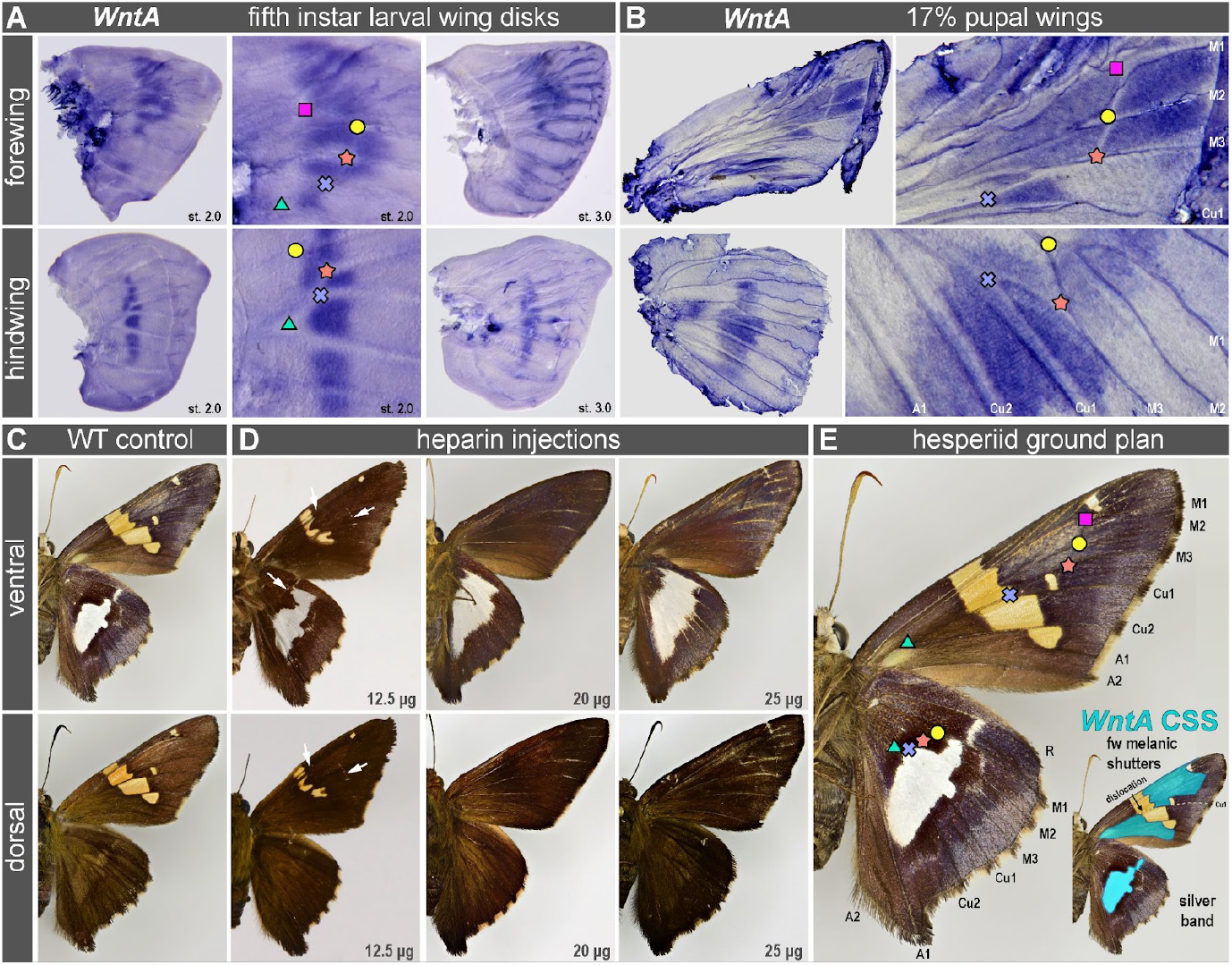
*WntA* marks CSS patterns with phenotypically dual effects in *E. clarus*. Landmarks denote the point of bifurcation of presumptive veins (here, tracheal tube lumens) as in Fig. 1A. **A**. *WntA* mRNA *in situ* hybridization in fifth instar wing disks. Stages 2.0 and 3.0 are based on nymphalid equivalent stages (Reed et al., 2007). **B**. *WntA* mRNA *in situ* hybridization in 17% pupal wings. **C**. Adult patterns in a wild-type control individual. **D**. Heparin-induced pattern modifications, including the reduction and loss of the forewing orange band, and the expansion of the hindwing silver band. Arrows: areas of pattern expansion (melanic “shutter” expansions in forewings, silver band expansion in hindwings) visible after intermediate doses of heparin injection. **E**. Landmarking and inferred projection of *WntA* expression on adult patterns, marking a dual CSS homolog that corresponds to melanic “shutters” in the forewing that flank the orange band, and to the silver band in the hindwing (see text for details).

### Transcriptomics of proximo-distal patterning in early pupal wings

The expression of *WntA* in the presumptive CSS pre-pattern implies that upstream developmental factors are partitioning the wing into proximo-distal subdivisions. To gain preliminary insights on this process, we conducted an RNAseq analysis of 12% pupal forewings and hindwings, each sectioned into proximal, medial, and distal portions (**Fig. 4A**). DESeq2 analysis identified 1,480 differentially expressed genes (DEGs) between the forewing and hindwing tissues (**Fig. 4B-C**): 931 with a forewing enrichment, and 549 genes with a hindwing enrichment including the Hox gene *Ultrabithorax* (*Ubx*), required for providing hindwing identity (Matsuoka and Monteiro, 2022; Tendolkar et al., 2021; Tendolkar et al., 2024; Van Belleghem et al., 2023; Weatherbee et al., 1998). The high-expression and over-representation of genes assigned to the extracellular matrix in the forewing transcriptome suggests that this tissue is more actively involved in chitin production than the hindwings at the 12% stage (**Fig. S3**).

**Figure 4.**
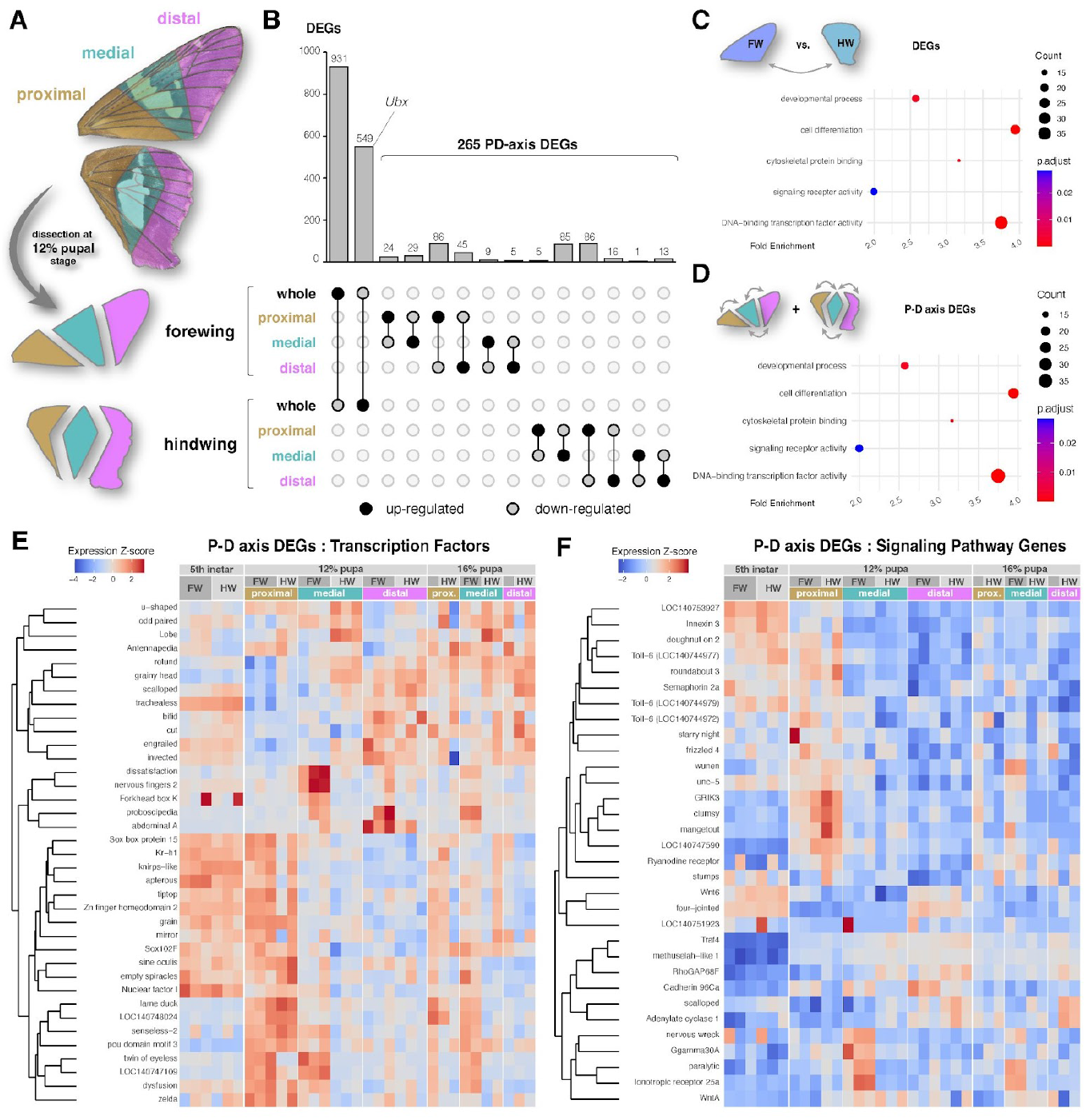
RNAseq and Differential Gene Expression analyses in *E. clarus* early pupal wings. **A**. Dissection of pupal wings along three proximo-distal sections, as projected on adult wings (top panel) to visualize the relative position of color pattern. Three biological replicates of each section were sequenced at the 12% pupal stage, except for proximal hindwings (two replicates). **B**. Upset plot featuring the results of pairwise comparisons of differential expression in 12% pupal wings between the forewing and hindwing, and all pairwise comparisons between sections within forewings and hindwings. The latter define a set of 265 proximo-distal axis DEGs. **C**. GO Enrichment Analysis for reduced GO categories, featuring 5 GO categories over-represented in the FW vs. HW DEGs. **D**. The same analysis identifies 4 GO categories over-represented in the P-D axis DEGs. **E-F**. Heatmap profiling of the 37 P-D axis DEGs classified as Transcription Factors (E), and signaling related genes (F). Columns each feature a different RNAseq sample. Forewing and hindwing wing disk samples dissected at the fifth instar larval stage, as well as 16% pupal wing samples sectioned as in panel A, are included to feature the expression of 12% stage DEGs over a wider temporal window.

Next, we investigated the set of genes showing a pattern of differential expression along the proximo-distal axis of each wing (P-D axis DEGs). A total of 265 P-D axis DEGs were identified (**Fig. 4B**), here as well we saw an over-representation of transcription factors and signaling pathway genes in GO enrichment analyses (**Fig. 4D**); this finding supports the idea that the regionalization of the wing epithelium involves spatial regulation of genes with a regulatory developmental function. To visualize their spatio-temporal expression patterns, we generated heatmaps for all 265 genes (**Fig. S2**) and for the subsets corresponding to transcription factors and signaling pathway genes as determined by GO term annotation (**Fig. 4E–F**), using RNAseq data from whole fifth instar wing discs and from proximal, medial, and distal sections of wings at the 12% and 16% pupal stages.

Expression profiling validates the enrichment of *WntA* expression in the medial region of the pupal hindwings where the silver stripe is patterned (**Fig. 5A**). As *u-shaped* (*ush*), *odd-paired* (*opa*), *lobe* (*L*) are up-regulated in the early medial hindwing in a similar fashion, it will be interesting to test in the future the expression of these transcription factors is connected to WntA signaling in presumptive color patterns. The transcription factors *homothorax* (*hth*), *tiptop* (*tio*, corresponding to *tio/tsh* in *Drosophila*), *zinc finger homeodomain 2* (*zfh2*) show proximal expression bias (**Fig. 5B**), consistent with previous transcriptomic studies of *Heliconius* early pupal wings (Hanly et al., 2019; Hines et al., 2012), and with their established roles as specifiers of proximal regions in *Drosophila* wing disks (Ruiz-Losada et al., 2018; Terriente et al., 2008; Zirin and Mann, 2004). Inversely, the transcription factor genes *rotund* (*rn*), *vestigial* (*vg*), *scalloped* (*sd*), *bifid/optomotor-blind* (*omb*), as well as the genes *Fat* and *Four-jointed* showed distal enrichment profiles (**Fig. 5C**).

**Figure 5.**
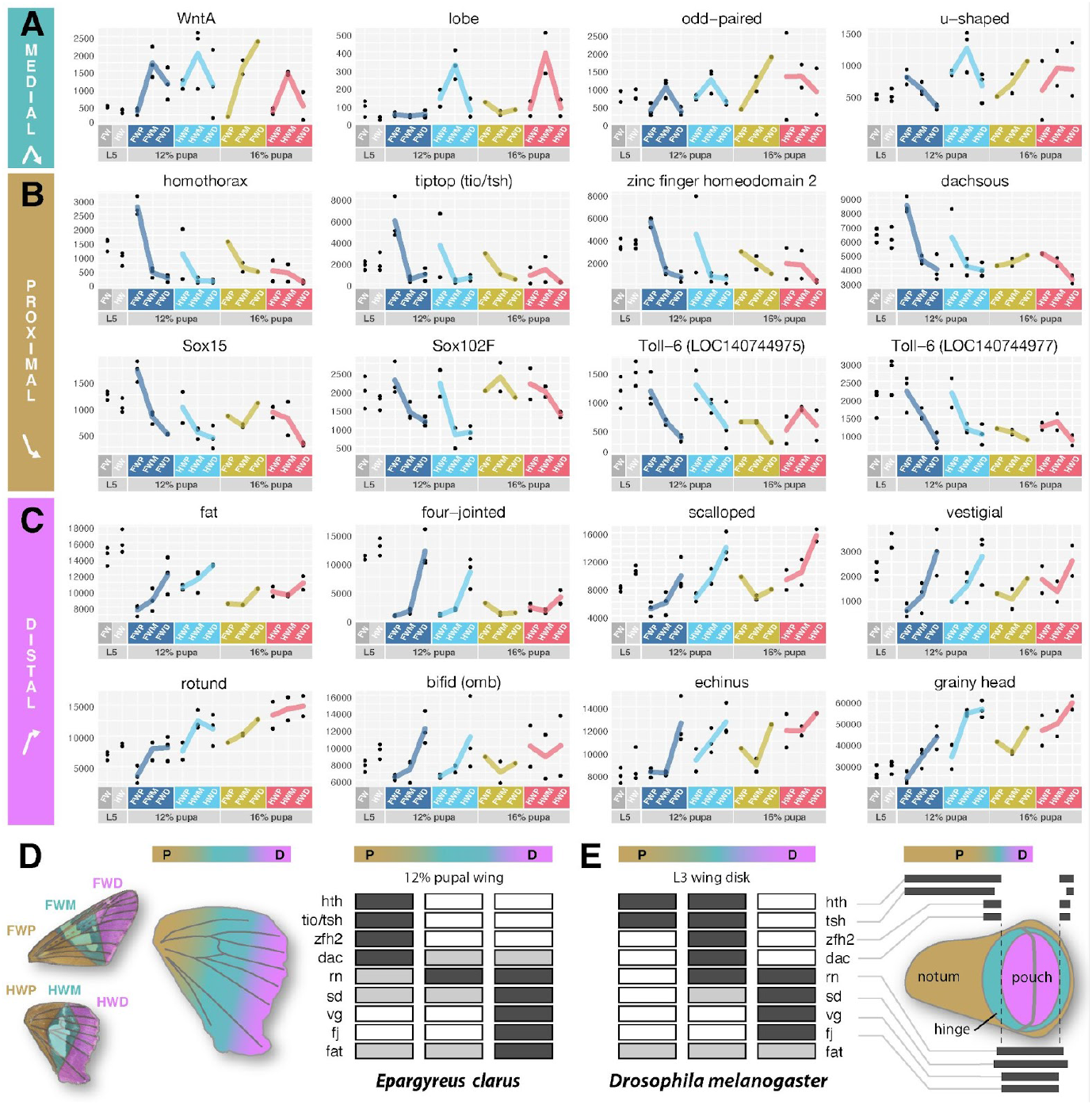
RNA-seq profiling of selected patterning genes. **A-C**. Gene expression profiles for select DEGs, with normalized count values plotted on the y-axis. Panels A, B and C each group genes by expression profile in the 12% hindwing. **A**. Genes with enrichments in the median region, similar to *WntA*: *lobe, odd-paired, u-shaped*. **B**. Genes with a proximal bias: *homothorax, tiptop, dachsous, zinc finger homeodomain 2* (*zfh2*), *Sox15, Sox102F*, and two tandem copies of *Toll-6* (LOC140744975 and LOC140744977). **C**. Genes with a distal bias: *fat, four-jointed, rotund, vestigial, echinus, grainy head, scalloped, bifid* (*omb*). **D-E**. Summary of the gene expression profiles of candidate proximo-distal specifiers in 12% *E. clarus* pupal wings (D), and compared to the expression of their orthologues in the *D. melanogaster* wing imaginal disk (E), as described in previous publications (Cho and Irvine, 2004; Everetts et al., 2021).

## Discussion

### How to identify CSS patterns in Hesperiidae?

Weaving relationships of homology between rapidly evolving patterns requires the identification of core mechanisms that have been maintained since the divergence of two lineages. The conserved expression of *WntA* in the medial region of the developing wings of nymphalids and hesperiids likely indicates homology of the CSS patterns between these two lineages that separated about 95 MYA (**Fig. 6A**). Schwanwitcsh, through his lifelong work examining pattern variation, had correctly predicted the position of the CSS and its dual nature in *P. sidae* (**Fig. 1D**). Indeed, *WntA* expression and heparin injections in *E. clarus* validated several properties of the CSS that are counterintuitive: first, that the forewing CSS is dislocated and acting as a melanic shutter, *i.e*. framing the more narrow stripes and dots of lighter color (**Fig. 6E**); second that the CSS can match a pattern of a different color identity than in the forewing, namely a white pattern in *P. sidae* and a silver band in *E. clarus*.

**Figure 6.**
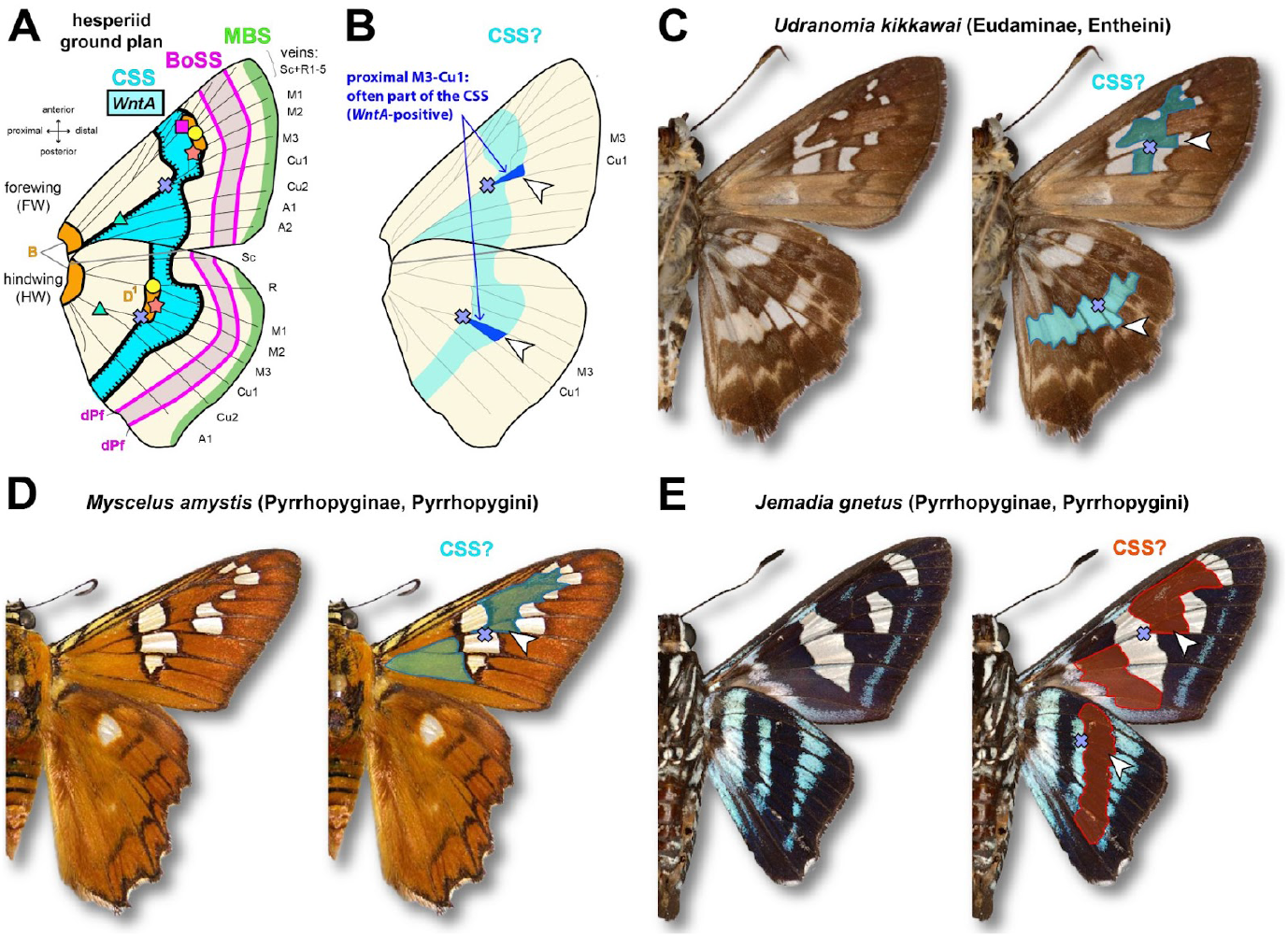
Extrapolation of the hesperiid ground plan to other species. **A**. Suggested hesperiid groudplan. Abbreviations and landmarks as in Fig. 1A. **B**. Based on empirical observations in nymphalids, the pattern element (arrowhead) immediately distal to the M3-Cu1 vein junction (cross) can often be used to infer the identity of the CSS. **C**. CSS inference in *Udranomia kikkawai*, a species from the same sub-family as *E. clarus* (Eudaminae), where a dual CSS is likely. **D-E**. Examples of inferred CSS patterns in the rapidly diverging Pyrrohopyginae subfamily of firetip skippers. The forewings of these two species feature possible cases of ‘pierellization’, with a dislocation of the CSS along the Cu1 vein, and with its distal boundary aligning with a basal patterning system in its anterior section. *Photo Credits*: Nick V. Grishin (C, E) and Daniel H. Janzen (D), retrieved from the Butterflies of America online repository (https://www.butterfliesofamerica.com).

These data highlight the evolutionary malleability of hesperiid wing patterns and generate testable hypotheses about CSS homology in this lineage. While the identification of this element remains challenging without visualizing *WntA* expression, we can propose initial clues on the identity of the CSS in species with medial patterns. In nearly all nymphalid and hesperiid species we assessed, *WntA* consistently marks the tissue immediately distal of the Cu1-M3 vein junction—which is landmarked as a blue cross here, and as a blue dot in our previous studies (Concha et al., 2019; Gallant et al., 2014; Hanly et al., 2023; Huber et al., 2015; Martin and Reed, 2014b; Mazo-Vargas et al., 2017b). As a rule of thumb, the proximal Cu1-M3 area (arrowheads in **Figs. 6B-6C**) is a good indicator of the color output of the CSS in a given wing surface, and we provide putative examples of this principle in a few Hesperiidae. Based on this, we propose that forewings of the rapidly diverging Pyrrohopyginae subfamily show a complete dislocation of the CSS along Cu1 (**Fig. 6D-6E**). This phenomenon would be analogous to the pierellization of many nymphalids (Nijhout, 1991; Otaki, 2021; Penz, 2017; Schwanwitsch, 1925), a term coined to describe the dislocation of the CSS and the coincidence of its distal boundary with a more proximal patterning system. Thus, the developmental mechanisms that fuel and constrain the evolution of wing patterns are similar between nymphalids, hesperiids, and beyond.

We also caution these principles should not be applied too dogmatically given that the dynamics of wing patterning remain poorly understood: as known exceptions among nymphalids, the Monarch *Danaus plexippus* and Gulf Fritillary *Agraulis incarnata* (formerly *Agraulis vanillae*) both underwent drastic shifts from the archetype of *WntA* expression as an antero-posterior medial stripe (Mazo-Vargas et al., 2017b; Mazo-Vargas et al., 2022b). The breathtaking diversity and rapid radiations of hesperiid wing patterns, particularly in contexts of wing mimicry and convergence (Li et al., 2019; Zhang et al., 2019), suggest that similar principles are at play in skippers and that other patterning tools than *WntA* may fashion this complex morphospace.

### Deep similarities between skipper and *Drosophila* P-D patterning

RNAseq profiling or early pupal wing development across three P-D subdivisions confirmed that *homothorax* and *tio/tsh* mark proximal regions of the lepidopteran wing (Hanly et al., 2019; Hines et al., 2012), and this expression is reminiscent of their role in specifying proximal identity in the *Drosophila* wing disk (Zirin and Mann, 2004). Interestingly, *dachsous* showed a proximal pattern that is inverted compared to the distal-enrichment patterns of *fat* and *four-jointed*. Dachsous and Fat are interacting protocadherins whose opposing gradients, modulated by the kinase Four-jointed, establish the Fat-Dachsous planar cell polarity pathway that orients tissue polarity (Thomas and Strutt, 2012). In *Drosophila*, this system also regulates the growth of larval wing disks, with *Dachsous* acting proximally in the body wall region, and Fat/Four-jointed acting more distally in the wing pouch, in conjunction with the Vestigial and Scalloped transcriptional co-factors (Cho and Irvine, 2004; Halder et al., 1998). Our finding that these genes are likewise enriched proximally (*dachsous*) or distally (*fat, four-jointed, vestigial, scalloped*) in skipper wings indicates that a P-D patterning module is deeply conserved. In spite of their distinct wing imaginal disks, this reinforces the idea that P-D patterning in both flies and butterflies relies on a conserved set of transcription factors and signaling modules (**Fig. 5D-5E**).

### New insights in the regulatory genetics of pupal wing development

While the development of wing tissues may follow conserved principles, the color patterns that decorate them also involved the evolution of novel gene expression, such as the pattern-specific domains of expression of *WntA* in Nymphalidae (Banerjee et al., 2023; Concha et al., 2019; Martin and Reed, 2014b; Mazo-Vargas et al., 2017b; Mazo-Vargas et al., 2022b), Papilionidae, Papilionidae (Mazo-Vargas et al., 2024), and in Hesperiidae. Little is known so far about the expression of *u-shaped, odd-paired*, and *Lobe* in lepidopteran wings, and further work is required to determine the extent to which they are associated with WntA patterning, and if their expression undergoes divergent or conserved evolution.

In addition to these medial signals (**Fig. 5A**), we also found enrichment profiles in the proximal region of the early pupal wing that were more pronounced, for the transcription factors *Sox102F, Sox15*, and tandem-paralogs of the Toll-6/Tollo receptor (**Fig. 5B**). Proximal enrichment was more pronounced in 12% wings, a stage the scale organ precursor cell lineage (SOP) has differentiated from other epithelial cells, and are progressively undergoing a second round of cell division to give rise to scale and socket daughter cells (Loh et al., 2025). As *Sox102F* marks epithelial (non-SOP) cells (Loh et al., 2025b), and as *Sox15* marks socket cells (Loh et al., 2025b; Loh et al., 2025b; Prakash et al., 2024), their proximal enrichment in our 12% RNAseq samples could be thus due to a wave-like event of cell differentiation starting in the proximal region, as suggested by time-lapse observations of the early pupal wing epithelium (Iwata et al., 2014; Loh et al., 2025). The function of *Toll-6* genes remains to be determined, possibly hinting at a novel role of Toll-like receptor signaling during lepidopteran wing patterning or cell specification.

## Methods

### Skipper butterflies

Immature stages of *Epargyreus clarus* (Cramer, 1775) were collected as larvae on kudzu vines around Washington, DC in the summers of 2018, 2019, and 2024. Larvae were provided kudzu cuttings fit into floral water vials in tupperware containers and reared in a growth chamber with 40-60% relative humidity and a 14:10 h light:dark cycle, set at 26°C or 32°C (see below). Pre-pupae were transferred to transparent cups and pupation time was recorded using a TX-164 timelapse camera (Technaxx).

### Genome sequencing and annotation

High-molecular weight DNA was extracted from an *E. clarus* female pupa using a Qiagen Genomic-tip 100/g before preparation and sequencing of a PacBio HiFi library on a PacBio Revio sequencer at the Institute for Genome Science at the University of Maryland. Sequencing reads (NCBI SRA SAMN38816873) were assembled into a primary haplotype assembly using *Hifiasm* with a variety of *-s* values (from the default value of 0.55 to 0.35). The best assembly was chosen based on the highest contiguity and lowest duplication rate (assessed with *Compleasm*). Despite testing a variety of *-s* values, duplicate contigs remained in the assembly and were subsequently removed using *purge_dups* (Cheng et al., 2021; Guan et al., 2020). Genome assembly quality was assessed using *BUSCO v.6.0.0* and *Compleasm* v0.2.6 using the *Lepidoptera_odb10* lineage dataset and *BlobToolKit* (Challis et al., 2020; Huang and Li, 2023; Tegenfeldt et al., 2025). Painting of Merian elements onto GCF_041222505.1 scaffolds (**Fig. 2C-D**) was performed with *Lep_BUSCO_Painter* v.1.0.0, using BUSCO ortholog assignments from the reference set of lepidopteran ancestral linkage groups (Wright, 2024; Wright et al., 2024).

Non-wing tissue transcriptomes generated for annotation purposes were obtained by tissue storage in TRI Reagent (Zymo Research), followed by RNAseq library preparation and sequencing at a target depth of 30M 150PE reads per sample by Azenta/Genewiz, and are available under the NCBI Bioproject PRJNA1165403. Genome annotation was performed by the NCBI Eukaryotic Genome Annotation Pipeline team; this resource is available as the *Epargyreus clarus WU_Ecla_fem_2.2* annotated reference genome (GCF_041222505.1). The GTF file produced from this annotation effort was manually curated for the purposes of this study to include the distant exon 1 of the gene *Cortex* from wing transcriptomic evidence generated from this study (**Data File S1**). Additionally, the gene feature list was manually curated to include functional annotations, such as the *Drosophila melanogaster* genes with sequence similarity found by performing reciprocal BLASTp between the *WU_Ecla_fem_2.2* RefSeq_protein sequences and the translated protein sequences from the FB2025_02 FlyBase release of the *D. melanogaster* genome annotation using the BLAST+ tool (Camacho et al., 2009). The column *gene_label* was also added to the gene feature list to apply recognizable and shortened gene names from the NCBI and *D. melanogaster* functional annotations for clear data visualization. Lastly, annotation names for the Wnt ligand genes were manually assigned using a phylogenetic analysis (**Fig. S1**), based on a maximum likelihood phylogeny computed using W-IQ-tree default parameters, from an amino-acid sequence alignment produced by MAFFT in Geneious Prime and trimmed with and filtered with Guidance2 with a reliability threshold of 0.101 (Katoh and Standley, 2013; Sela et al., 2015), which added predicted *E. clarus* Wnt proteins to a previously published set of Wnt protein sequences (Hanly et al., 2021).

### In situ hybridizations

An antisense riboprobe targeting *E. clarus WntA* (NCBI Genbank: XM_073090400.1) was PCR amplified from larval wing disk cDNA using the following primers (Forward: 5’-CGAAGCAGCATTCGTACACG-3’; *T7-promoter*+Reverse: 5’-*TAATACGACTCACTATAGGG*GGTAGCCTCTTCCACAGCAT-3’), transcribed with a Roche T7 DIG RNA labeling kit, purified with Ambion MEGAClea r columns, and stored at −80°C. Detection of *WntA* mRNA expression in larval and pupal wings followed previously published procedures (Hanly et al., 2023; Martin and Reed, 2014a), using 30 ng/mL of riboprobe during hybridization steps, and BM Purple (Roche) for the staining procedure. Pupal wings were dissected at 48-52 hr after pupation at 26°C, corresponding to about 17% of pupal development (average total pupal development time at 26°C was measured as 292 h, N =8). Images were taken using a Nikon D5300 camera mounted to a Nikon SMZ800N trinocular dissecting microscope, equipped with a P-Plan Apo 1X/WF 0.105 NA 70 mm objective and a Nikon C-DS stand for diascopic mirror illumination.

#### Heparin injections

Pupae between 3-14 hr after pupa formation (32°C) were injected in their abdomen with 2.5-5 μL of 5 μg/μL heparin sodium salt (Sigma-Aldrich, H3393) or with water, using microcapillary needles fit into a Nanoject III microinjector. All treated pupae emerged into adult butterflies, which were pinned and imaged with a Nikon D5300 camera mounted with a Micro-Nikkor 105mm f/2.8G lens.

### Pupal wing RNAseq

Wing transcriptomes are available under the NCBI Bioproject PRJNA660444 and were prepared as follows, using a published procedure (Hanly et al., 2019). Larvae and pupae were reared at 26°C until dissection and storage in RNA later. Pupal wings were dissected at 36 hr (12 % pupal development) and 48 hr after pupation in cold PBS (16%), and cut using microdissection scissors into six compartments: FWP (proximal forewing), FWM (medial forewing), FWD (distal forewing), HWP (proximal hindwing), HWM (medial hindwing), and HWD (distal hindwing) using the developing veins as landmarks (**Fig. 4A**). For RNA extraction, wing tissues were transferred into 2 mL tubes containing 500 μL of Trizol Reagent (Thermo Fischer), and homogenized with a Tissue Lyser for 2 min at 30 Hz. 200 μL of chloroform were added to the lyzate before vigorous shaking for 15 seconds, settling for 3 minutes at room temperature, and centrifugation at 13,000 rpm for 15 minutes at 4 °C. The aqueous top phase was transferred to a new tube, measured, and the same volume of 70 % EtOH was added drop by drop to avoid localized precipitation. Total RNA was purified using the RNeasy kit (Qiagen), treated with DNAse I (Ambion) at 37 °C for 10 minutes before addition of 5 μL of inactivation reagent, and stored at −80 °C. Library preparation, and sequencing were performed by the Sequencing and Genomics Facility (University of Puerto Rico Rio Piedras, San Juan, Puerto Rico). Samples were sequenced at either 75 SE or 150 PE on an Illumina NextSeq sequencer. Adapters were removed using Trimmomatic v0.39 using default parameters with the exception of the values SLIDINGWINDOW:4:20 and MINLEN:85 specified for PE samples (Bolger et al., 2014). Post-trimming, FastQC v.0.11.8 was used to verify all adapters were successfully removed and to assess read quality (Andrews, 2025). Reads were aligned to the *Epargyreus clarus WU_Ecla_fem_2.2* reference genome (GCF_041222505.1) with STAR v.2.7.11 (Dobin and Gingeras, 2016). STAR alignment was repeated with the IntronMotif output from the original alignment to better resolve splice junctions prior to read counting.

### Differential Gene Expression Analyses

*FeatureCounts* (Subread v.2.0.8) was used to perform read counting (Liao et al., 2014) with the manually curated GTF annotation file (**Data File S1**). This count dataset was used to perform differential expression analyses using DESeq2 in RStudio (Love et al., 2014). First, we investigated which genes were differentially expressed between forewings and hindwings at 12% pupal development, by collapsing the counts from each compartment (proximal, medial, and distal) of each individual forewing or hindwing prior to count normalization. The experimental design ~ *wing_type* was then used to perform differential expression analysis with a significance threshold of adjusted *p* < 0.05. The R package *ggplot2* (Wickham and Sievert, 2009) was used to produce Volcano and MA plots to visualize these data (**Fig. S3**).

Next, to define the set of P-D axis DEGs at 12% pupal development, we used the experimental design *~ compartment*, resulting in 6 combinations of pairwise comparisons (3 between the 3 forewings compartments, and 3 between the 3 hindwing compartments) with a significance threshold of adjusted *p* < 0.05. To build a heatmap of z-score expression of genes across compartments and developmental timepoints, the experimental design *~ compartment* was re-run on the entire count dataset to produce normalized counts for all samples, and used to build gene expression profile plots for genes of interest and a heatmap restricted to the set of P-D axis DEGs identified at the 12% pupal developmental stage using the R package *ggplot2* (Wickham and Sievert, 2009). Hierarchical clustering was applied to the dataset to cluster by similar gene expression profiles prior to visualization by performing variance stabilizing transformation using the function *vst()* from DESeq2, z-score scaling, and use of the R package *ggdendro* (Vries and Ripley, 2024). The clustering *data* and dendrogram were then incorporated into the heatmap visualization produced by *ggplot2*.

Gene Ontology (GO) Enrichment Analyses were performed separately on both the wing_type and compartment DEG datasets with *clusterProfiler* (Xu et al., 2024), using the *E. clarus* GO annotation hosted on the NCBI FTP server (*WU_Ecla_fem_2.2*), after it was mapped to the *slimGO_agr* and *slimGO_Drosophila* standardized subsets using *Map2Slim* in the OWLTools v4.5.29 package (Mungall et al., 2024). Dot plots for each GO enrichment analysis were produced by *ggplot2* (**Fig 4C-4D; Fig S3**). Transcription Factors were then defined from *slimGO_agr* re-mapped annotations as the subset of genes matching the Molecular Function categories GO:0003700 or GO:0008134 (“DNA-binding transcription factor activity”, or “transcription factor binding”). Signaling pathway genes were similarly defined as the subset of genes with a Biological Process annotation matching GO:0023052, GO:0038023, or GO:0005102 (“signaling”, “signaling receptor activity”, or “signaling receptor binding”). Filtered heatmaps with the corresponding gene category subsets with hierarchical clustering of gene expression profiles were then produced according to the aforementioned procedure (**Fig 3E-3F**).

## Supporting information

Data S1: Manually curated GTF Genome Annotation File

Data S2: Manually curated Gene Annotation List

## Acknowledgements

We thank Martha Weiss, Allison Brackley, Grace Jeschke, Anna Ren, and John Lill for their support rearing *E. clarus* larvae, Ling Sheng Loh for help dissecting pupal wings, Yadira Ortíz for RNAseq library preparations, Anyi Mazo-Vargas for providing a DNA extraction, and Daniel Davis for assisting with genome sequence submission. The annotation of the *E. clarus* genome was graciously carried out by the NCBI Eukaryotic Genome Annotation Pipeline team. We are also grateful for the Butterflies of America initiative to have made available an important photographic record of New World Hesperidae.

## Data and resource availability

The *WU_Ecla_fem_2.2* NCBI RefSeq genome sequence and annotation for E. clarus is available on NCBI Datasets under the identifier GCF_041222505.1. The manually curated annotation GTF file and gene list derived from this resource can be found as **Data File S1** and **Data File S2**, respectively. Wing transcriptomes are available on the NCBI SRA under NCBI Bioproject PRJNA660444. The detailed results from all differential expression analyses performed in this study can be found in **Tables S1-S11**. All code associated with the differential expression analyses performed in this study can be found in a GitHub repository (Alqassar, 2025).

## Supplementary Information

**Data S1**: Manually curated GTF Genome Annotation File

**Data S2**: Manually curated Gene Annotation List

**Figure S1**: Assignment of *E. clarus* Wnt ligand family genes to eight orthology groups

**Figure S2**: Heatmap of all 265 P-D axis DEGs

**Figure S3**: Significance, expression level, and GO enrichment analyses of differential gene expression between the 12% pupal forewings and hindwings

**Table S1**: Table containing the DESeq2 normalized counts for all genes in the genome produced during the forewing vs hindwing (wing-type) analysis

**Table S2**: Table containing the DESeq2 Results from the forewing vs hindwing (wing-type) analysis

**Table S3**: Table containing the DESeq2 normalized counts for all genes in the genome produced during the wing compartment analysis at 12% pupal development

**Table S4**: Table containing the DESeq2 Results from the forewing proximal vs forewing medial compartment analysis at 12% pupal development

**Table S5**: Table containing the DESeq2 Results from the forewing medial vs forewing distal compartment analysis at 12% pupal development

**Table S6**: Table containing the DESeq2 Results from the forewing proximal vs forewing distal compartment analysis at 12% pupal development

**Table S7**: Table containing the DESeq2 Results from the hindwing proximal vs hindwing medial compartment analysis at 12% pupal development

**Table S8**: Table containing the DESeq2 Results from the hindwing medial vs hindwing distal compartment analysis at 12% pupal development

**Table S9**: Table containing the DESeq2 Results from the hindwing proximal vs hindwing distal compartment analysis at 12% pupal development

**Table S10**: Table containing the DESeq2 normalized counts for all genes in the genome per wing compartment analysis for all developmental timepoints

**Table S11**: Table containing the data plotted in the full heatmap in Figure S2

**Figure S1.**
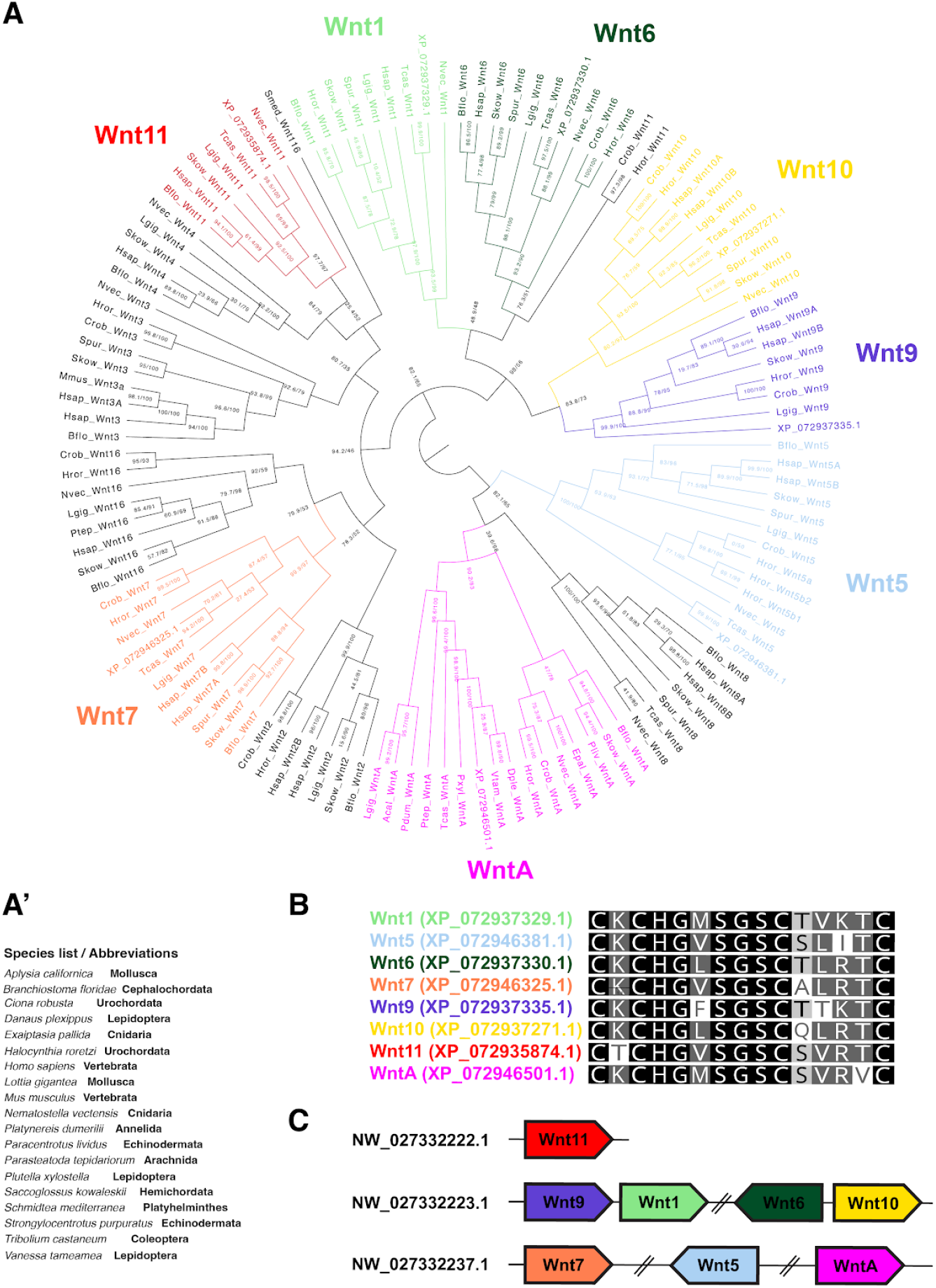
Assignment of *E. clarus* Wnt ligand family genes to eight orthology groups. **A**. Maximum likelihood reconstruction of *E. clarus* and reference Wnt proteins (Hanly et al., 2021). Branch support is indicated by SH-aLRT % values / ultrafast bootstrap % values. **A’**. Species abbreviation key for the phylogeny. **B**. Amino-acid alignment of the eight *E. clarus* Wnt thumb regions. **C**. Arrangement of Wnt genes within the *E. clarus WU_Ecla_fem_2.2* genome assembly.

**Figure S2.**
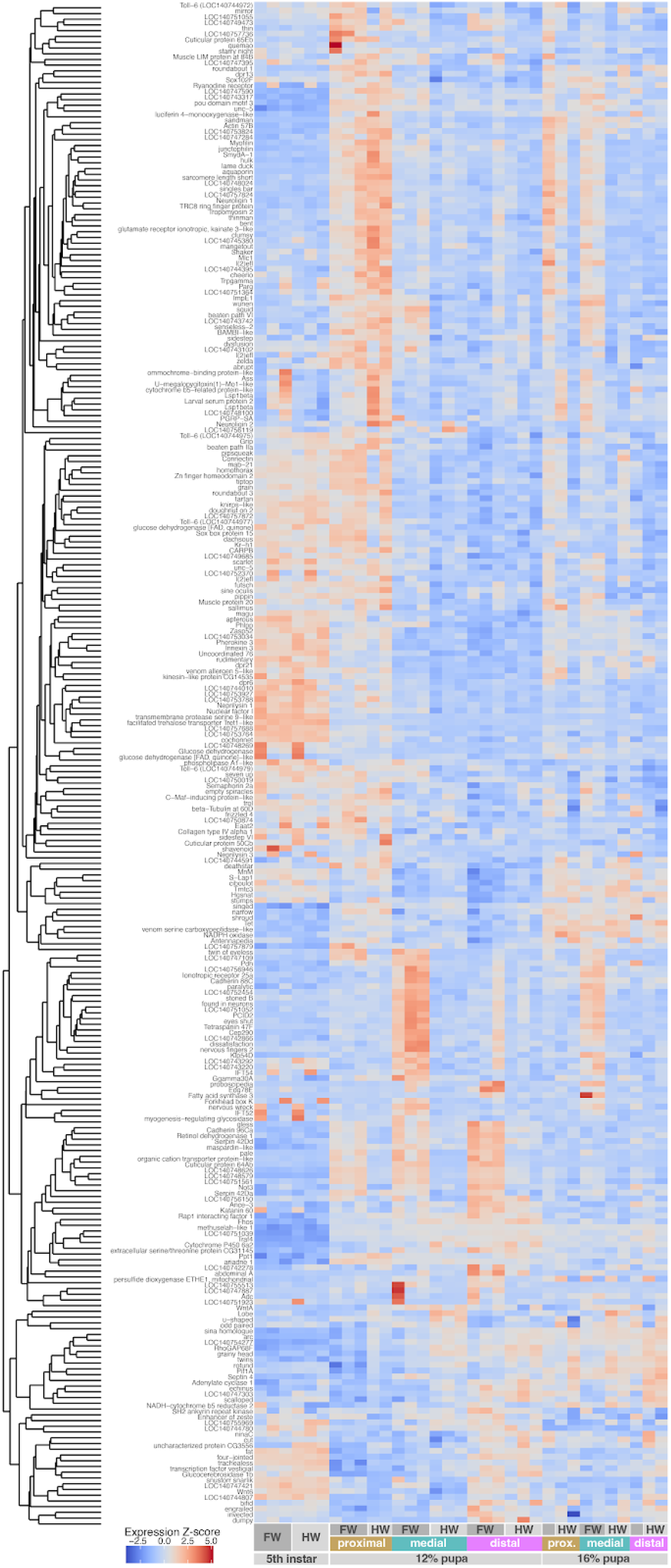
Heatmap of all 265 P-D axis DEGs. Heatmap of all 265 proximo-distal axis differentially expressed genes, color-coded using a z-score normalization of their expression levels across all samples and time points. A dendrogram reflects the hierarchical clustering of genes by similar expression profiles.

**Figure S3.**
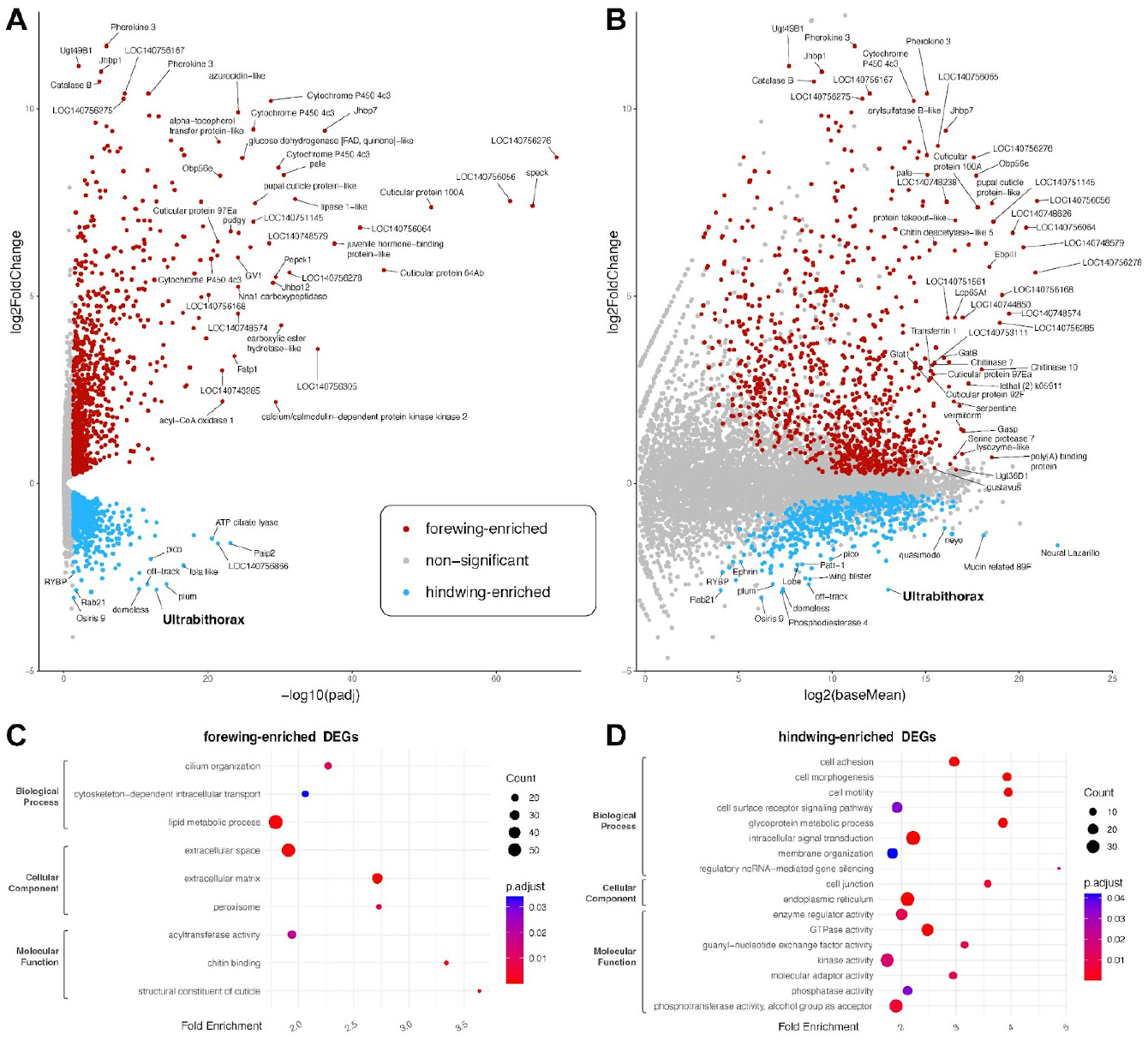
Significance, expression level, and GO enrichment analyses of differential gene expression between the 12% pupal forewings and hindwings. Transcriptomes appear asymmetric between the forewing and hindwing. This is partly due to the high expression of forewing-specific genes, many of which with annotations associated with cuticle development, suggesting this reflects a role of the forewing in the maturation of cuticle, perhaps relating to the juxtaposition of the forewing and pupal case. **A**. Volcano plot of 12% pupae forewing vs hindwing DEGs. The y-axis is Log2FoldChange Expression and the x-axis is −log10(adjusted p-value). **B**. MA-plot visualization of the same genes. The y-axis is log2FoldChange Expression, and the x-axis is the log2 of average normalized expression (baseMean in DEseq2). **C**. Results of a GO enrichment analysis of forewing-enriched DEGs, using the *slimGO_Drosophila* standardized subset. Enriched categories are clustered by ontological classes (Biological Process, Cellular Component, and Molecular Function). **D**. The same analysis performed for hindwing-enriched DEGs.

